# Segmentation-Enhanced CycleGAN

**DOI:** 10.1101/548081

**Authors:** Michał Januszewski, Viren Jain

## Abstract

Algorithmic reconstruction of neurons from volume electron microscopy data traditionally requires training machine learning models on dataset-specific ground truth annotations that are expensive and tedious to acquire. We enhanced the training procedure of an unsupervised image-to-image translation method with additional components derived from an automated neuron segmentation approach. We show that this method, Segmentation-Enhanced CycleGAN (SECGAN), enables near perfect reconstruction accuracy on a benchmark connectomics segmentation dataset despite operating in a “zero-shot” setting in which the segmentation model was trained using only volumetric labels from a different dataset and imaging method. By reducing or eliminating the need for novel ground truth annotations, SECGANs alleviate one of the main practical burdens involved in pursuing automated reconstruction of volume electron microscopy data.

## Introduction

Volume electron microscopy (VEM) methods have enabled nanometer-resolution imaging of biological tissue over fields of view that now routinely span millions of cubic microns^1^. A primary application of VEM is the determination of neural circuit connectivity, in which detailed reconstructions of individual neurons are combined with image-based identification of synapses ^2,3^. Due to the infeasibility of manually performing reconstruction of thousands of neurons ^4^, automation of this task has emerged as a major practical requirement for revealing neural circuits in VEM datasets ^5^.

Significant progress in automated neuron reconstruction has been achieved through the development of machine-learning (ML) based image segmentation methods optimized for tracing of neurites in VEM data ^6–10^. ML methods require training data that has traditionally been created for each specific acquisition context, which differ by choice of species, brain area, imaging method, staining protocol, and other sample-specific details. In one recent study, 408 hours of human time was invested to manually annotate 131 million voxels of songbird brain that was then used to train an ML-based segmentation approach ^6^. While the trained algorithm reconstructed songbird neurons with high accuracy, the prospect of repeatedly performing hundreds of hours of manual annotation for each such study is clearly undesirable ^11^.

A classical approach to improving the generalization abilities of an ML system is dataset augmentation, in which domain-specific modes of variation are computationally “simulated” and applied to a database of training examples ^12^. This approach has been used to great effect for improving *intra*-dataset generalization performance of VEM segmentation methods by introducing synthetic forms of rotation and reflection ^13^, elastic deformation ^14^, and image defects such as misalignment, missing sections, and out-of-focus sections ^7^.

Such data augmentation strategies have not, however, succeeded in enabling strong *inter-*dataset generalization. Therefore we pursued a new approach, Segmentation-Enhanced CycleGAN^15^ (SECGAN), in which we learn a model that “translates” raw image content between two different VEM datasets. By translate we mean we render the images from one dataset in the style of another so that it resembles the shapes and textures of the other dataset while retaining its original cellular boundaries. If one dataset lacks the volumetric ground truth necessary to train a segmentation algorithm, we can segment it by translating its raw data and then applying a segmentation algorithm trained on the other dataset for which we have ground truth (Fig. 1a). Note that this approach does not require any notion of correspondence or alignment between images in the two datasets; the translation function is learned in an unsupervised fashion.

**Figure 1.**
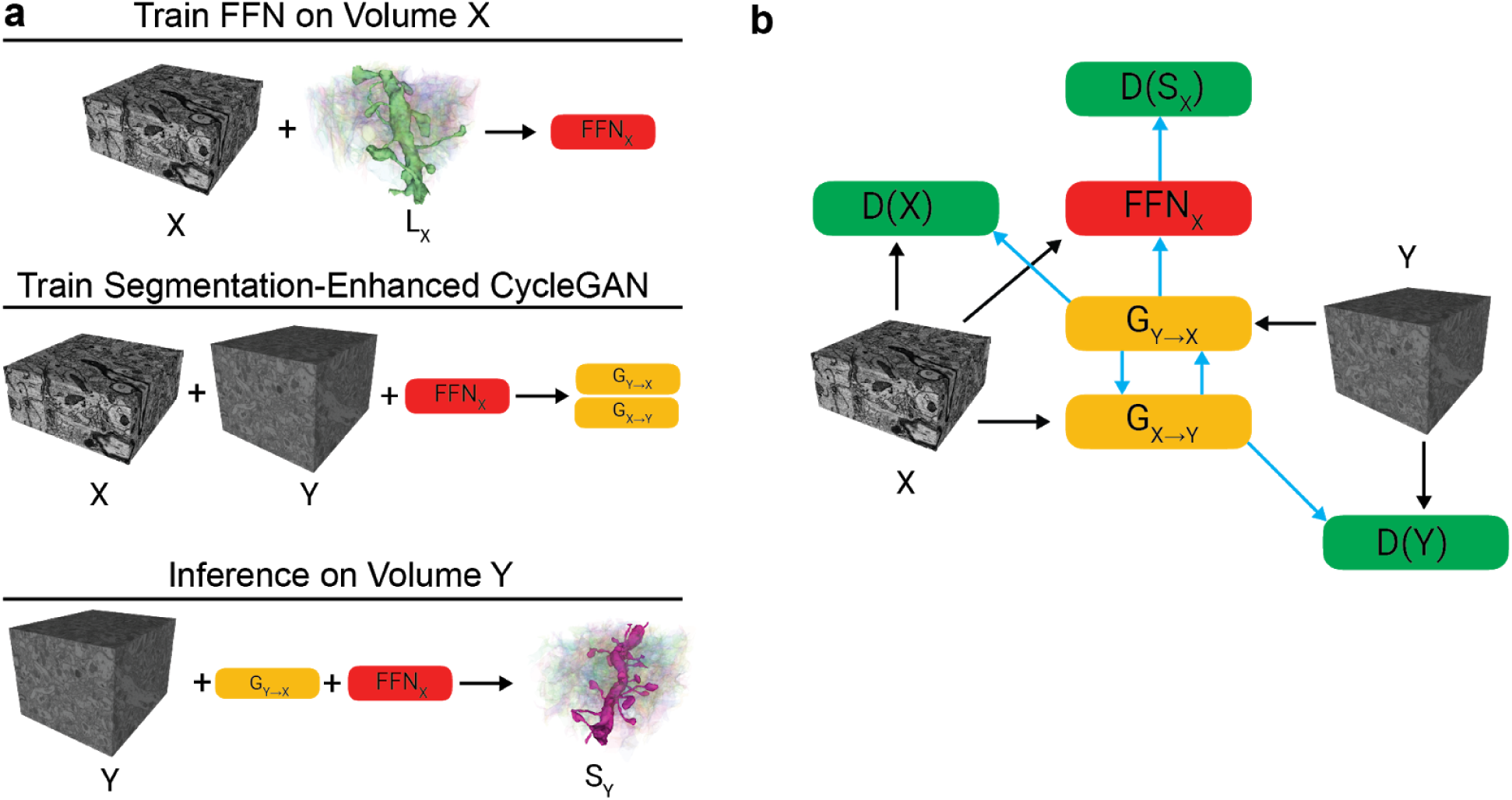
Segmentation-Enhanced CycleGAN. (a) Schematic of overall training and inference workflow. A flood-filling network (FFN) is trained on ground truth data associated with volume *X*. The trained FFN is then used to segment *Y* by translating data from *Y* into *X* using the SECGAN. (b) Components and information flow of Segmentation-Enhanced CycleGAN training, which consists of three discriminator networks, two generator networks, and an FFN. Blue arrows indicate pathways along which there is also reverse flow of gradients.

## Results

The core of the SECGAN approach is a cycle-consistent generative adversarial network (CycleGAN), which is a neural network that bidirectionally translates data between two domains ^15^. In our case, a “generator” function G_X→Y_ takes input data from VEM dataset *X* and is trained to produce outputs that are visually similar to a target VEM dataset *Y*; the training of G_X→Y_ principally relies on backpropagation signals from a “discriminator” function D(*Y*) which predicts whether its inputs are true samples from *Y* or “fake” samples produced by G_X→Y_ ^16^. A reverse generator G_Y→X_ and discriminator D(*X*) are trained in a similar fashion. A “cycle consistency loss” is applied that composes the generators and optimizes G_Y→X_(G_X→Y_(*x*)) ≈ *x* for samples from *X* and G_X→Y_(G_Y→X_(*y*)) ≈ *y* for samples from *Y*. Each generator must therefore embed enough information about the source domain in its output in order for the other generator to be able to approximately recover the original input. A key property of the CycleGAN is that its training is fully unsupervised and does not rely on any notion of a per-sample correspondence between *X* and *Y*.

SECGAN enhances a CycleGAN with additional components that bias the generator G_Y→X_ toward producing images that yield plausible automated neuron segmentation results. Specifically, samples from *X* and the output of G_Y→X_ are provided as input to a flood-filling network (FFN), which is a neural network architecture designed for 3d neuron segmentation ^6^. The FFN is pre-trained on volumetric ground truth from *X* and its parameters are unchanged during SECGAN training. A third discriminator D(*S*_*X*_) receives the output of the FFN and tries to predict whether those segmentations were generated by data sampled from *X* or G_Y→X_ (Fig. 1b). D(*S*_*X*_) is trained simultaneously with the other components of the SECGAN and thus provides an additional source of backpropagation signals that influence parameters of G_Y→X_ and G_X→Y_.

We evaluated the SECGAN approach by performing automated reconstruction of a mouse somatosensory cortex sample (“SegEM”) imaged using serial block-face scanning electron microscopy (SBEM) at a voxel size of 11×11×28nm ^10^. In a control condition, “dedicated,” we trained an FFN using ground truth located within the SegEM dataset itself, which resulted in excellent segmentation accuracy (Fig. 2 and Fig. 3). We then segmented the SBEM dataset using a model trained on a somatosensory cortex dataset (“SNEMI3d”) imaged using a different VEM method, automated tape-collecting ultramicrotome scanning electron microscopy (ATUM-SEM), at a voxel size of 6×6×29nm ^17^. Without SECGAN transfer of the VEM data (condition “CLAHE”), the FFN trained on SNEMI3d reconstructed the SegEM dataset poorly; *with* SECGAN transfer, the results were in fact slightly superior to the dedicated model, despite the FFN operating in a “zero-shot” transfer mode ^18^ in which no training was performed on labeled data from the target volume (note that checkpoint selection used skeleton metrics on a subvolume of the target VEM data, see Methods). Standard CycleGAN transfer was also worse than SECGAN transfer (see Fig. 3).

**Figure 2.**
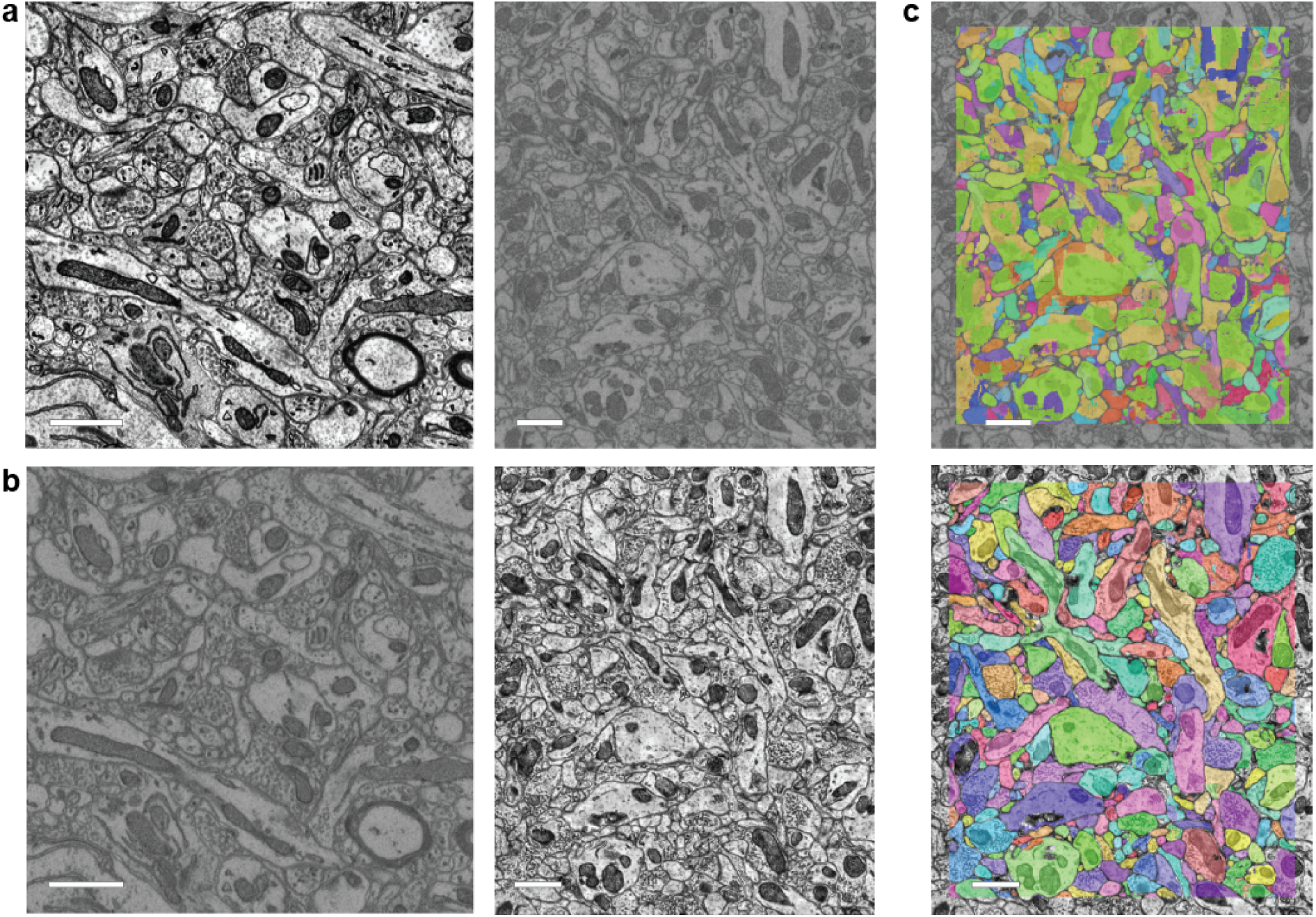
Qualitative analysis of results. (a) Raw (*x-y*) data from SNEMI (left) and SegEM (right) VEM data. (b) Bidirectional translations given by CycleGAN: SNEMI→SegEM (left) and SegEM→SNEMI (right). (c) SegEM segmentation results using FFN trained on SNEMI ground truth: CLAHE-only baseline (top) versus SegEM→SNEMI SECGAN translation (bottom). Scale bar 1 μm.

**Figure 3.**
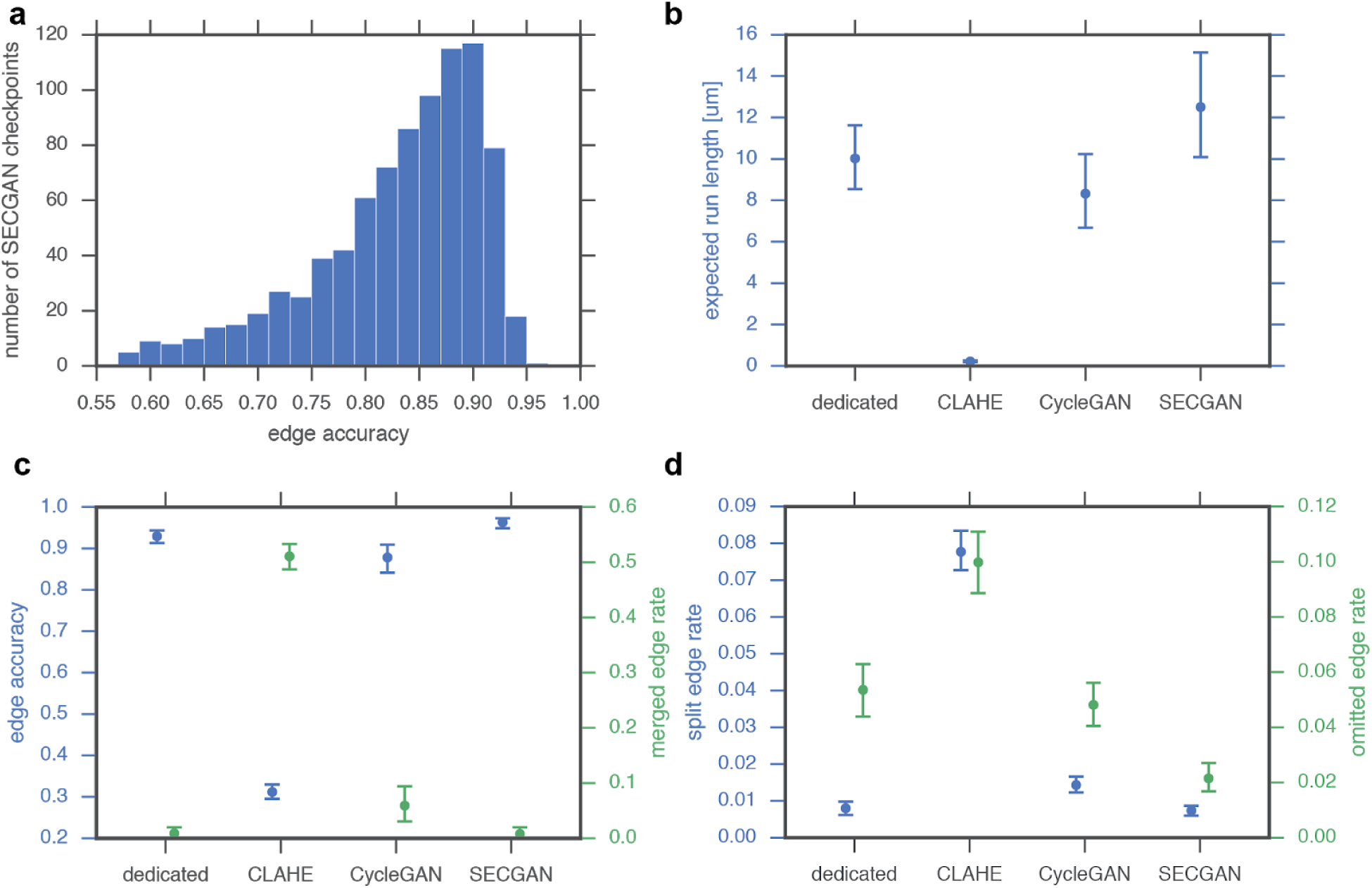
Quantitative analysis of SECGAN and baseline results. (a) Number of SECGAN checkpoints that lead to transfer results with a particular segmentation (edge) accuracy, as derived from a single training run. (b-d) Expected run length and skeleton accuracy metrics of SegEM segmentation results generated by SECGAN-transfer and CycleGAN-transfer versus CLAHE-baseline and a dedicated (non-transfer) model trained directly on SegEM volumetric ground truth.

## Discussion

SECGANs alter raw VEM data in order to enable model transfer. Therefore it is reasonable to wonder whether the alterations are safe; could using a SECGAN lead to incorrect, misleading, or biased analyses? In Figure 4, we highlight specific aspects of the SegEM → SNEMI translation that the SECGAN performed. Based on the quantitative and qualitative results in Fig. 2-4, as well as the fundamentally local nature of SECGAN processing, we believe there is little reason to suspect that the transfer procedure leads to globally biased neuron reconstructions (segmentation of even a single neuron relies on the accumulation of predictions over millions or billions of voxel locations). Though in our experiments we found the ultrastructure to be well preserved by the transfer, the use of methods such as SECGAN for more local analyses of VEM data, such as the detection of synapses, needs to be more carefully evaluated. Related work has found that using neural networks to significantly alter raw microscopy data can be useful for the purposes of super-resolution ^19,20^, alignment and interpolation ^21^, content restoration ^22^, and inferring “virtual” fluorescent labels in the absence of physical antibody stains ^23^.

**Figure 4.**
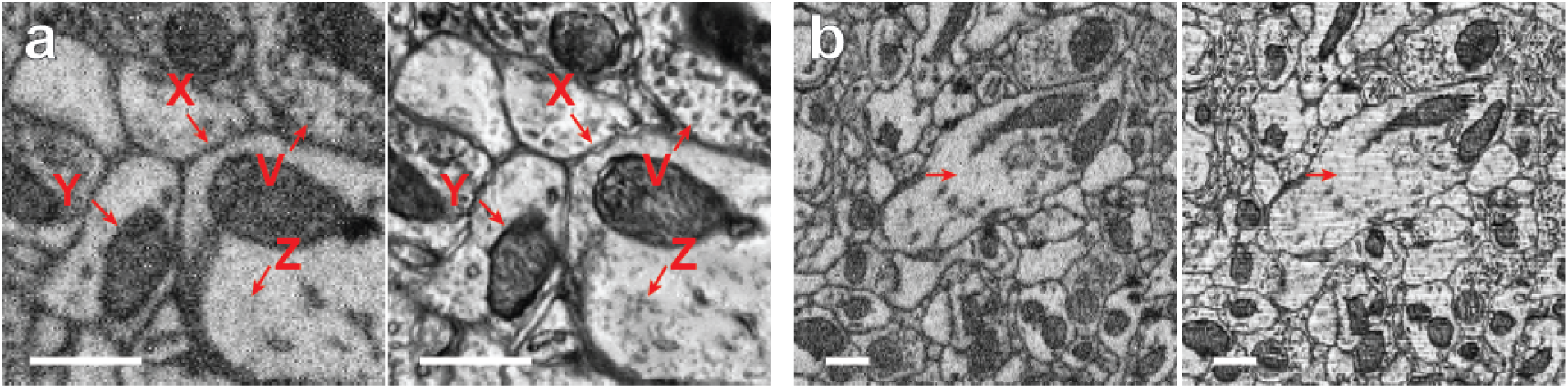
Selected examples of SECGAN results. (a) In-section (*x-y*) view of SegEM data post-processed with CLAHE (left) versus output from a SegEM → SNEMI SECGAN model (right). Note: (X) altered space between membranes, (Y) altered internal structures in the mitochondrion, (Z) altered texture inside the neurite, (V) similar number and position of vesicles. (b) Cross-section (*x-z*) view. Alignment of the SECGAN-processed data is unchanged. SECGAN reproduced section-to-section intensity differences present in the target (SNEMI3d) image, visible here as horizontal stripes. Scale bar 0.5 μm.

Dataset augmentation is an important approach to improving the generalization capabilities of an ML system. However, augmentation is fundamentally less appropriate when addressing modes of variation that are difficult to “a priori” describe algorithmically, such as the idiosyncratic, high-dimensional, and locally correlated set of pixel intensity differences that may arise from imaging brain tissue using different microscopy methods or different staining protocols. Moreover, it is sensible to take advantage of the raw data itself when modeling the translation from one dataset to another and in situations where unlabeled raw data is copiously available, as is generally the case when dealing with reconstruction and analysis of VEM data. Hence SECGANs offer a powerful tool for transferring reconstruction models among VEM data and can significantly reduce the practical burden associated with successful connectomic reconstruction of diverse datasets.

## Acknowledgements

We thank Jeff Lichtman and Jörgen Kornfeld for helpful comments on the manuscript.

## Online methods

### SECGAN training

The SECGAN system is built using three neural network architectures: the generator (G), the discriminator (D) and the segmenter (S), as outlined in Fig. 1b. In addition to the D(*X*) and D(*Y*) discriminators which use image data as input similarly to the CycleGAN, we introduce the segmentation discriminator D(S), which processes single-object segmentation data only. We also experimented with combining the D(*X*) and D(S) into a single discriminator with a two-channel (raw image, single-object segmentation) input, and removing D(*X*), both of which resulted in worse segmentation quality.

G and D consume and produce images rescaled to [-1, 1] as (original / 127.5 - 1.0), where we assume 8-bit unsigned image intensities in the original images. The input to S is rescaled as (original - 128) / 33, in accordance with ^6^. The output of S has the form of a probability map and is scaled as (probability - 0.5) when it becomes the input of D(S).

The SECGAN training procedure has 4 steps, in which G_X→Y_, D(*S*)+D(*X*), G_Y→X_, and D(*Y*) are sequentially optimized. The generator weights are optimized by minimizing the following loss:

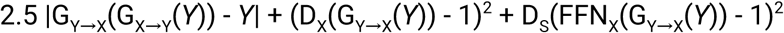

where mean absolute difference is used for the cycle-consistency loss (first term), and mean squared difference is used for the discriminator losses, and where the loss values are averaged over the examples in the batch and over voxels within every example. The discriminator weights are optimized by minimizing the following loss:

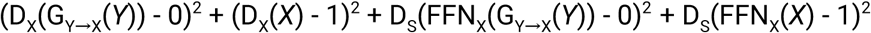

The segmenter weights are held fixed during training of the SECGAN.

We implemented the SECGAN system in TensorFlow and trained it with asynchronous stochastic gradient descent on 8 NVIDIA V100 GPUs, with a batch size of 8 and learning rate of 0.0001. Training examples for the unsupervised training of the SECGAN were sampled randomly from SNEMI3d testing and training volumes (both 512 x 512 x 100 voxels), and from SegEM cortex testing (401 x 401 x 351 voxels) and training (740 x 868 x 251 voxels) volumes.

### Neural network architectures

We used a 3d generalization of ResNet-18 ^24^ as the architecture for all three discriminator networks, and a stack of 8 residual convolutional modules as the architecture of the generator networks. We also tested a U-net-style generator, as well as a simple convolution-pooling network as the discriminator, and found these alternatives resulted in worse segmentation quality.

In the generator, each residual module consists of two 3d convolutions with (3, 3, 3) kernels, 32 feature maps, operating in VALID mode, with the ReLU activation function used for first convolution and linear activation used for the second convolution. The residual skip connections used center-cropping of the source to match the target volume size. The output of the last residual module was passed through a pointwise convolution layer with a single feature map and *tanh* activation function to form the generated image.

The architecture of the discriminator followed that of ResNet-18, but we eliminated the final classification layer and the global average pooling preceding it, and used the output of the last residual module directly, which we found necessary for the network to train stably.

For segmentation, we used a flood-filling network with the architecture and training parameters as reported in ^6^ but trained on 2x in-plane downsampled SNEMI3d challenge data (512 x 512 x 100 labeled voxels; voxel size: 12×12×29 nm; see Supplementary for a discussion of the impact of voxel size discrepancies on the transfer process).

### Intensity inversion

We observed that the CycleGAN and SECGAN models sometimes settle into a regime where a call to each generator network inverts the intensity of the input image (bright regions become dark, and vice versa). The generator loss consists of two terms -- the cycle-consistency loss, and the discriminator loss. An intensity-inverting pair of generators can have a low value of the first term, but not of the second one if the discriminator is trained enough to distinguish whether an image is real or generated.

Empirically, the intensity inversion depends on the random initial state of the network, becomes apparent after a few hundred training iterations, and the network does not recover from it during training, even as the discriminator performance improves in time. This may be related to the higher weight (2.5) used for the cycle-consistency loss, which we however found necessary for the network to reach a state where it produces useful results.

To work around the intensity inversion problem, at training steps 300 to 2000, for every training example we identified the lowest (*y*_*l*_) and highest (*y*_*h*_) intensity voxel in the *X->Y* generated image, and checked whether in the full-cycle image (*X->Y->X*) the intensities of these two voxels were reversed G_Y→X_(*y*_*l*_) > G_Y→X_(*y*_*h*_). If so, we restarted the training of the network by choosing a new set of random values for all its internal parameters.

### SegEM segmentation evaluation

To quantitatively test our segmentation accuracy, we used the “cortex training” subvolume released as part of the SegEM challenge together with manually traced skeletons (https://segem.rzg.mpg.de/webdav/SegEM_challenge/skeletonData/). A 640 x 768 x 201-voxel fragment of the subvolume (corresponding to 7.2 x 8.5 x 5.6 μm of tissue) was exhaustively traced so that each neurite visible in the volume has an associated skeleton.

A manual review of the skeletons revealed several topological inconsistencies (see supplementary for details), which we corrected. We then used the modified skeletons as ground truth for evaluation of our segmentation results with skeleton metrics^6^. To compensate for inaccurate placement of skeleton nodes and minor variations in the spatial extent of the segments, a merge error was counted only when a segment overlapped more than 2 nodes from each of two or more skeletons.

The 95% confidence intervals shown in Fig. 3 were computed with the bootstrap method, with 10,000 resamples from the set of ground truth skeletons.

### SECGAN checkpoint selection

We observed significant segmentation quality differences when the same FFN was used to segment images obtained with generators initialized with different snapshots of the SECGAN network weights (“checkpoints”). A typical distribution of edge accuracies is shown in Fig. 3a. After an initial training period lasting ∼15k steps of the SECGAN network training procedure where the accuracy clearly increased over time, we did not find any patterns in the time series of the evaluation results. We speculate that the varying performance is caused by the generator network oscillating between reproducing different features of the target image. For instance, we observed that the checkpoints for which the segmentation has the worst edge accuracy correspond to generated images without altered space between membranes of adjacent neurites (see Fig. 4a), which causes the FFN to incorrectly merge them together. Fig. 2c and Fig. 3b-d show data for the best segmentation result.

## Supplementary

### SegEM “cortex training” skeleton modifications

We manually reviewed the released skeleton tracings of the SegEM “cortex_training” subvolume, and decided to ignore the following skeleton merges from evaluation:

- 705,1: two parts of the same axon
- 8,402: dendritic spine and shaft
- 708,709,308: fragments of the same axon
- 661,597: dendritic spine and shaft, connected outside of the skeletonized subvolume
- 458,406: dendritic spine and shaft
- 475,580: skeletons trace the same neurite
- 510,692: two glial fragments, connected at 234, 682, 41
- 656,710: spine head and rest of the dendrite
- 397,214: connected around 336, 219, 29
- 680,109: 680 is an organelle (?)
- 398,707: dendrite and spine
- 604,704: dendrite and spine
- 16,399,669: dendrite and spine

### Resolution mismatch

In the experiments reported in the main text, the source and target datasets have different voxel sizes (6×6×29 nm for SNEMI3d, and 11×11×29 nm for SegEM) and therefore we used a 2x in-plane downsampled version of SNEMI3d (i.e., 12×12×29nm voxel size) in all experiments. We have also applied the SECGAN system to other VEM dataset pairs with larger voxel size differences, and from different species. Qualitatively, we observed that the quality of the segmentation of the target dataset degrades with larger physical discrepancies between the source and the target, particularly where the source has a lower voxel resolution than the target. In all cases, we found the results to be useful at least for the purpose of targeted ground truth generation through object-based proofreading for training of a dedicated FFN model, the details of which will be reported elsewhere.

### SNEMI→SegEM translation

To check the effectiveness of the translation procedure in the opposite direction, we trained a second SECGAN network using the SegEM-trained FFN used for the “dedicated” results in Fig. 3. We used this SECGAN to segment the SNEMI3d training volume, applying mirror padding in all directions in order to minimize edge effects and let the FFN create segments extending all the way to the border of the volume. We also skeletonized the SNEMI3d training volume with TEASAR and used the skeletons as ground truth to compute segmentation metrics. We postprocessed the skeletons to erode tips up to 200 nm and to reduce inter-node distance to an average of 300 nm in order to make them more similar to ones generated manually by human tracers.

The best segmentation obtained with the SECGAN reached 83.6% edge accuracy at 0.6% merge, 7.2% split, and 8.6% omitted edge rates. The segmentation looked qualitatively good, but the split and omitted edge rates were significantly higher than those obtained in the transfer process in the opposite direction (Fig. 3b-d). These errors appeared to be caused primarily by the smallest caliber processes, particularly when they were oriented parallel to the imaging plane.

